# A Transcriptional Lineage of the Early *C. elegans* Embryo

**DOI:** 10.1101/047746

**Authors:** Sophia C. Tintori, Erin Osborne Nishimura, Patrick Golden, Jason D. Lieb, Bob Goldstein

## Abstract

**HIGHLIGHTS:**

‒ RNA-seq on each cell of the early *C. elegans* embryo complements the known lineage
‒ We measured the zygotic activation specific to each unique cell of the embryo
‒ We identified genes that are functionally redundant and critical for development
‒ We created an interactive online data visualization tool for exploring the data

**eTOC BLURB:** *C. elegans* is a powerful model for development, with an invariant and completely described cell lineage. To enrich this resource, we performed single-cell RNA-seq on each cell of the embryo through the 16-cell stage. Zygotic genome activation is differential between cell types. We identified hundreds of candidates for partially redundant genes, and verified one such set as critical for development. We created an interactive online data visualization tool to invite others to explore our dataset.

**SUMMARY:** During embryonic development, cells must establish fates, morphologies and behaviors in coordination with one another to form a functional body. A prevalent hypothesis for how this coordination is achieved is that each cell’s fate and behavior is determined by a defined mixture of RNAs. Only recently has it become possible to measure the full suite of transcripts in a single cell. Here we quantify the abundance of every mRNA transcript in each cell of the *C. elegans* embryo up to the 16-cell stage. We describe spatially dynamic expression, quantify cell-specific differential activation of the zygotic genome, and identify critical developmental genes previously unappreciated because of their partial redundancy. We present an interactive data visualization tool that allows broad access to our dataset. This genome-wide single-cell map of mRNA abundance, alongside the well-studied life history and fates of each cell, describes at a cellular resolution the mRNA landscape that guides development.

## INTRODUCTION

An outstanding challenge of developmental biology is to explain how differential gene expression promotes the fundamental processes of embryonic development. Such processes include determining the fate of each cell, moving cells relative to each other to produce structures such as organs, and changing the composition and shape of each cell to perform metabolic or structural functions. Genomic approaches developed over the past decade have made it possible to generate comprehensive rosters of every transcript’s abundance in an organism or tissue during key developmental events. In this study, we have measured the abundances of all mRNAs in each cell of the early *C. elegans* embryo. In doing so, we have quantified the divergence of the genetic expression of these cells as they begin to perform diverse functions in the embryo.

The *C. elegans* embryo is a powerful and well-established system for studying cell biology and development (Figure 1A), and was chosen as a model organism in part because the entirety of development can be tracked with single-cell resolution (Sulston et al. 1983). The timing and orientation of every cell division, apoptotic event, and cell migration has been documented, but performing genomic studies with a matching resolution has been a challenge. Until recently, genomic protocols required collection of embryos in bulk, but *C. elegans* fertilization is staggered, rendering embryos asynchronous with each other. There is no practical system in place for culturing single cell types, leaving the only source of bulk biological material imprecisely staged samples that are usually composed of mixed cell types. Low-input RNA-sequencing (RNA-seq) methods developed within the last five years offer a solution to the genomics problem; a single *C. elegans* cell can be precisely identified and defined both in space and time.

**Figure 1:**
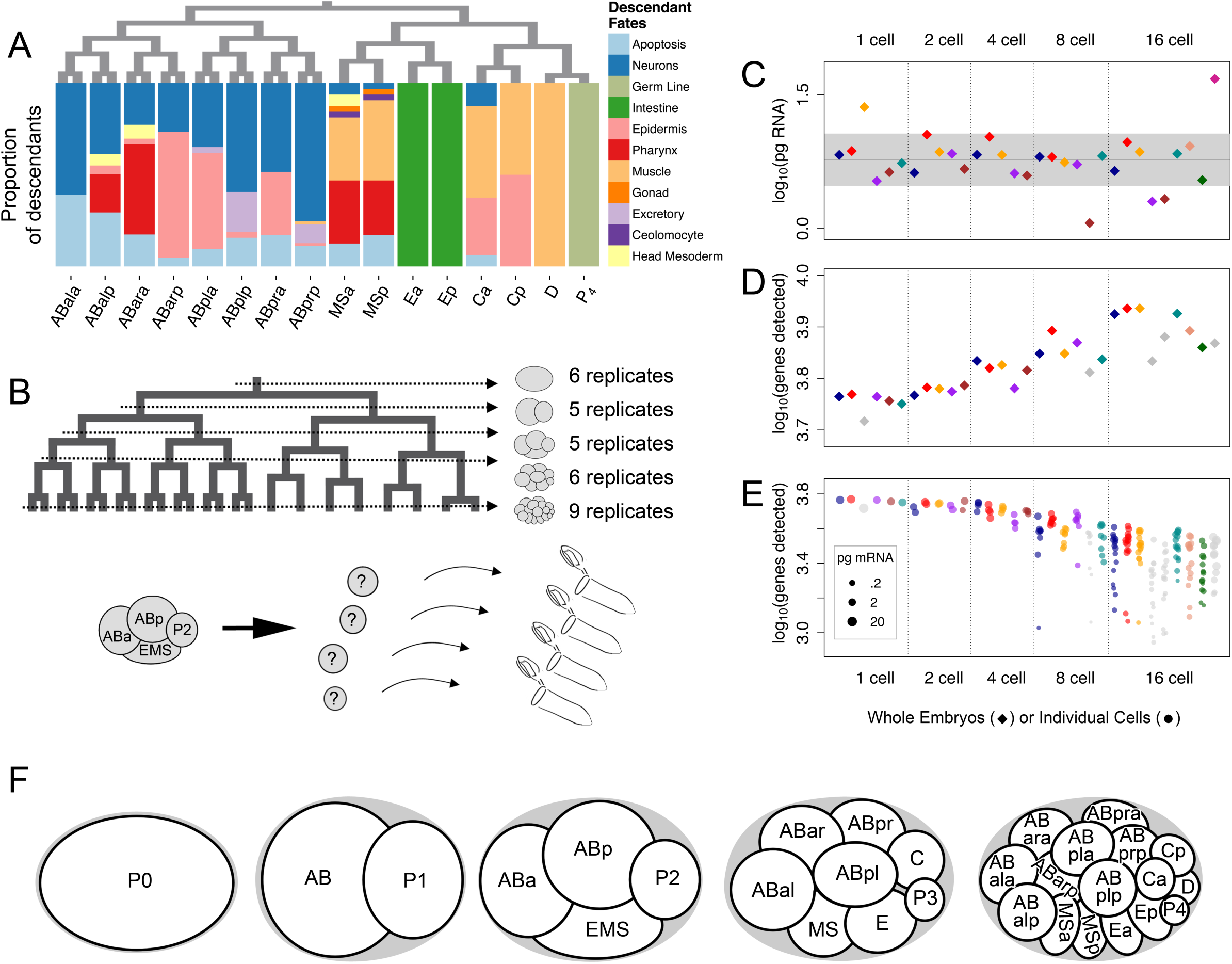
Single-cell mRNA-seq libraries for complete sets of cells from *C. elegans* embryos of the 1-, 2-, 4-, 8- and 16-cell stages. (A) Terminal cell fates of descendants of each cell of the 16-cell embryo. Terminal fates were calculated from Sulston et al. 1983, and refer to cell fates at the time of the first larval hatching. (B) Schematic of samples that were hand-dissected and prepared for scRNA-seq. 4-cell stage is diagrammed below for illustration. (C) The total mass of mRNA detected from each embryo (diamonds). Embryos whose total mass of mRNA differed from the average by more than one standard deviation (plotted outside of gray band) were excluded from subsequent analyses. (D) The number of genes whose transcripts were detected in each whole embryo (diamonds). (E) The number of genes whose transcripts were detected in each individual cell (circle). (F) Key of the names of each cell from the zygote to the 16-cell stage.

Understanding the full suite of mRNAs expressed in the *C. elegans* embryo has long been of interest. Whole-embryo mRNA timecourses revealed that thousands of genes are dynamically regulated at these early stages (Baugh et al. 2003; Baugh et al. 2005). Aided by advances in low-input RNA-seq technology of the last few years, researchers have interrogated the transcriptomes of the embryo by manually dissecting cells and performing RNA-seq. Due to the difficulty of identifying cells once they are dissected, only the 2-cell stage embryo has been sequenced at an entirely single-cell resolution (Hashimshony et al. 2012; Hashimshony et al. 2015; Osborne Nishimura et al. 2015). One study has performed transcript profiling of some single cells and some clusters of cells from later stages (Hashimshony et al. 2015). In this study we have sequenced each cell of an individual embryo in replicate for embryos up to the 16-cell stage. We used a combination of hand-dissections of each cell of a single embryo, and a unique strategy for identifying the dissected cells.

Many of the interesting phenomena of early development are transcriptionally regulated in *C. elegans*, including morphogenesis and cell fate specification (Edgar et al. 1994; Sommermann et al. 2010; Broitman-Maduro et al. 2006). Much of what we know about the genetics of these events has been gleaned from traditional genetic screens, which have a blind spot for pleiotropic genes and genes with partially redundant functions (Wieschaus 1997; Sawyer et al. 2011). With high-throughput sequencing, we can identify the genes whose transcript abundances correlate with morphogenesis, differentiation, or other phenomena, regardless of a gene’s possible pleiotropy or redundancy.

Here we present a transcriptional lineage of early *C. elegans* development, a map of all transcripts in each cell through the first stages of development. We generated this map by performing single cell RNA-seq (scRNA-seq) on each cell from the zygote to the 16-cell stage. We address previously unanswered questions about the differential activation of the zygotic genome in each cell, describe spatially dynamic gene expression, and identify previously unknown partially redundant genes that are critical for development. To maximize the usefulness of our dataset to the scientific community, we introduce a novel, publicly available interactive data visualization tool.

## RESULTS

### Transcriptome Diversity Among Cells of the Embryo Increases Over Time

Each cell at each stage in the early *C. elegans* embryo has a name, a known life history and fate, and is identifiable by its position relative to other cells (Sulston et al. 1983). We performed scRNA-seq on manually-dissected, individual cells from 1-, 2-, 4-, 8- and 16-cell stage embryos, with a minimum of 5 replicates for each sample (Figure 1B). We note that due to asynchronous cell divisions there is no true 16-cell stage, but we use this term for convenience (details in Experimental Methods). We sequenced the mRNA of each cell separately, knowing which embryo the cell came from but not knowing its identity, and used its transcript profile to identify its cell type *post hoc*. Cell size and cell division timing gave us some clues of the identities of 19 of the 31 cell types. For example, all the anterior (AB descendant) cells at the 8- and 16-cell stages divide in synchrony with each other, and the germ cell precursors at the 2- to 8-cell stages are considerably smaller than the rest of the cells (purple in Figure S1, and Extended Methods). These visual clues provided independent support for the results of our *post hoc* cell identity assignments (Figure S1 and described below).

In total, we generated 219 transcriptomes, describing quantitative expression levels for 8,575 detected genes (>25 RPKM). We aggregated data from cells of the same embryo to calculate whole-embryo statistics. To calculate the mass of mRNA in each cell, and thereby each whole embryo, we used spike-in controls from the External RNA Control Consortium (Baker et al. 2005). The mass of mRNA detected was relatively constant between stages, though embryos of later timepoints showed higher variability (Figure 1C). Among 31 whole embryos, five embryos had an mRNA mass more than one standard deviation above or below the average and were excluded from further analysis (details in Table S1). To evaluate changes in transcriptome complexity over time in both individual cells and whole-embryos, we calculated the number of mRNA species detected in each single-cell transcriptome and whole-embryo aggregation (Figure 1D,E). We noticed an increase in transcriptome complexity in whole embryos over time (>25 RPKM in any contributing cell), but a decrease in complexity in individual cells. The increase in whole-embryo complexity could be due to either cell-specific activation of the zygotic genome, or to the fact that a larger number of single-cell libraries constitute the whole embryo total at later stages, potentially allowing for fewer false negatives when compared to the small number of transcriptomes that make up whole-embryo values at earlier stages.

Before we could test the validity of the transcriptomes generated, we first needed to identify the cell type of origin for each transcriptome.

### Posterior Cells of the Embryo Have Distinct Cell-Specific Signatures Involving Hundreds of Genes

Many of the cell types we sampled are enriched for transcripts of one or a few known marker genes, which we were able to use to assign identities to our transcriptomes. A multi-gene clustering approach has been shown to be more effective at grouping replicates of a cell type than a single‐ or few-gene approach (Björklund et al. 2016; Jaitin et al. 2014; Grün et al. 2015; Satija et al. 2015). We used an iterative Principle Component Analysis (PCA) strategy (described below) to group transcriptomes by cell type, thereby collapsing our 219 transcriptomes down to 18 groups of identical or related cell types. We then used known marker genes to assign identities to each of these 18 groups (Figure 2). The 18 groups were P_0_, AB, P_1_, ABa, ABp, EMS, P_2_, ABxx (granddaughters of AB), MS, E, C, P_3_, ABxxx (great granddaughters of AB), MSx (daughters of MS), Ex (daughters of E), Cx (daughters of C), D and P_4_. Some of the groups that contained multiple cell types were later sorted into more specific groups (Figure 3).

**Figure 2:**
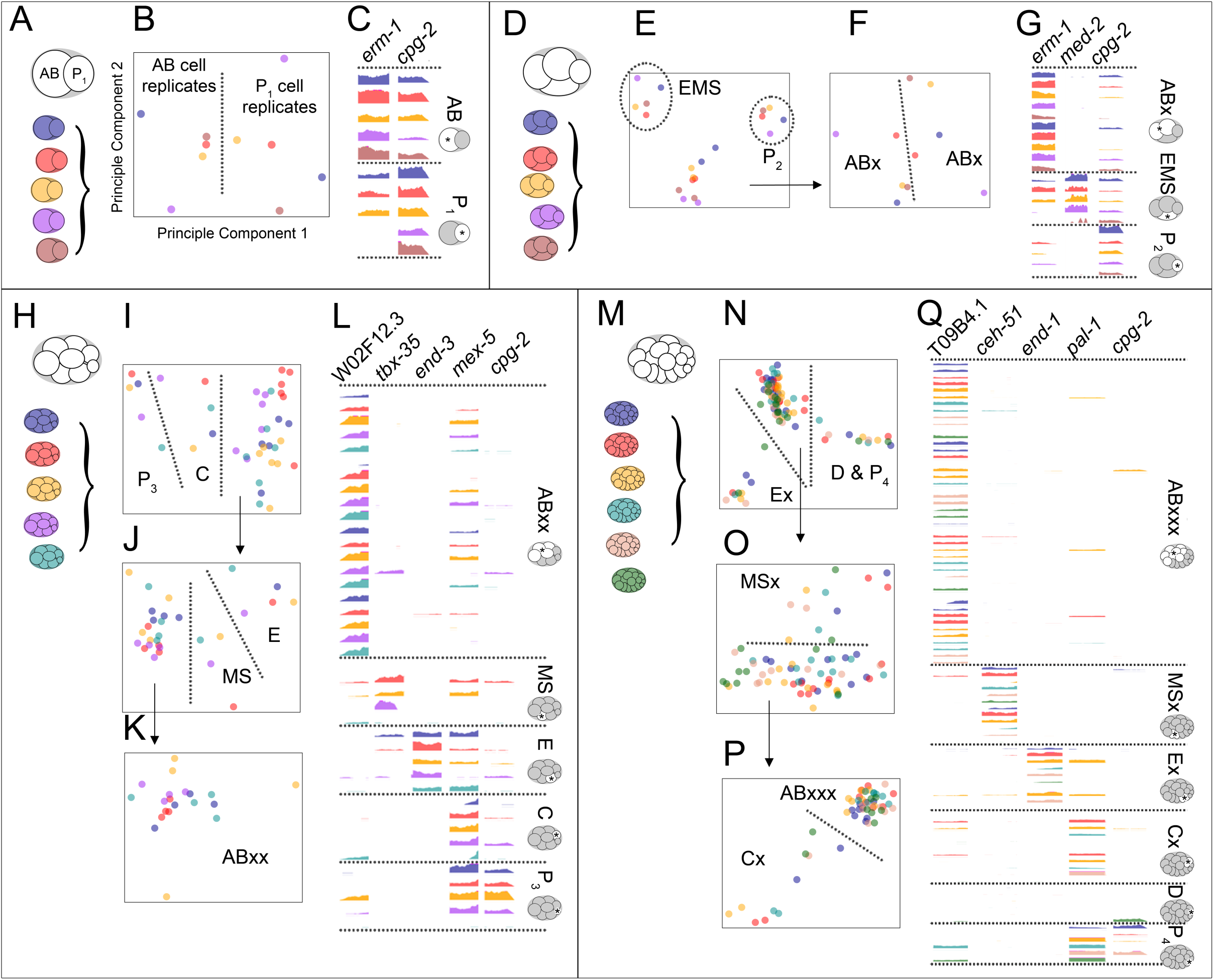
Replicates of each cell type were grouped by transcript signatures and identified by candidate gene expression. (A-C) Transcriptomes of cells from the 2-cell stage (A) were subjected to Principle Component Analysis (PCA) (B) using only data from reproducibly differentially enriched genes, as selected by our algorithm (details in Experimental Procedures). (C) Genome browser tracks of the last exon of *erm-1* (AB-enriched) and *cpg-2* (P_1_-enriched). Colors correspond to embryo of origin. Height of tracks indicate read count density. All y-axes of genome browser tracks are scaled consistently within each panel. (D-G) Transcriptomes of cells from the 4-cell stage (D) were subjected to PCA (E). (F) PCA of the 10 transcriptomes that were not resolved in (E). (G) Genome browser tracks of the last exon of *erm-1* (AB-enriched), *med-2* (EMS-enriched) and *cpg-2* (P_2_-enriched). (H-L) Transcriptomes of cells from the 8-cell stage (H) were subjected to PCA using iteratively generated sets of informative genes (I-K). (L) Genome browser tracks of the last exon of W02F12.3 (ABxx-enriched), *tbx-35* (MS-enriched), *end-3* (E-enriched), *mex-5* (C‐ and P_3_-enriched), and *cpg-2* (P_3_-enriched). (M-Q) Transcriptomes of cells from the 16-cell stage (M) were subjected to PCA using iteratively generated sets of informative genes (N-P). (Q) Genome browser tracks of the last exon of T09B4.1 (ABxxx-specific), *ceh-51* (MSx-specific), *end-1* (Ex-specific), *pal-1* (Cx‐ and P_4_-specific), and *cpg-2* (P_4_-specific).

**Figure 3:**
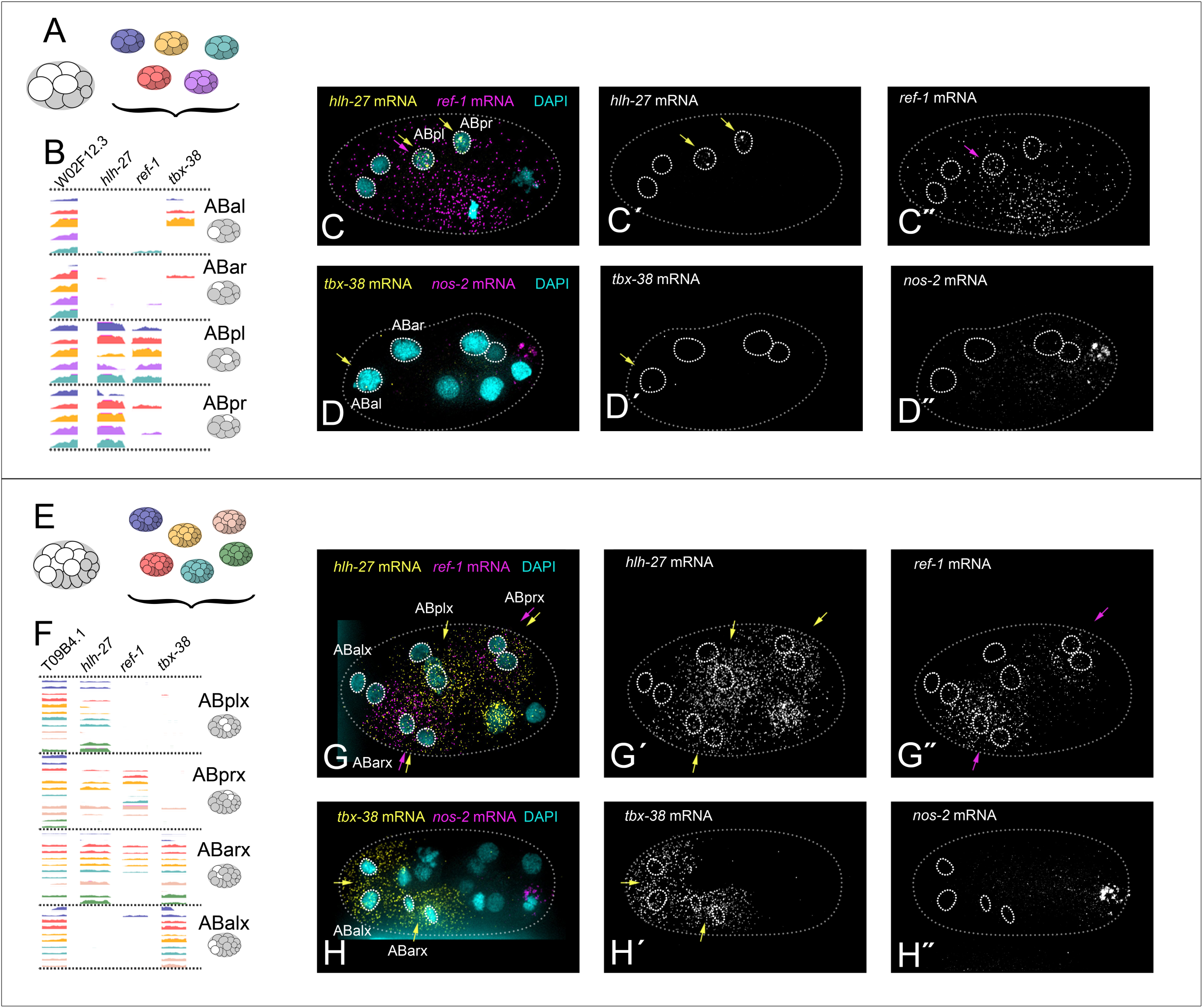
Differential transcript enrichment of notch target genes in cells that could not be distinguished by global transcript signatures. (A) AB descendants from five replicates of the 8-cell stage embryo. (B) Genome browser tracks of ABxx transcriptomes, sorted into groups based on expression of notch target genes *hlh-27*, *ref-1* and *tbx-38* (Extended Methods). Last exons only are shown. (C) Example of smFISH targeting *hlh-27* (C′, yellow arrows) and *ref-1* (C″, purple arrows) transcripts in intact 6‐ or 8-cell stage embryos (*hlh-27* pattern seen in 100% of embryos, n=4. *ref-1* pattern seen in 75% of embryos, n=4. Remaining embryo showed ubiquitous *ref-1* staining). (D) Example of smFISH targeting *tbx-38* (D′, yellow arrows) in intact 8-cell stage embryos (pattern seen in 33% of embryos, n=3. 67% of embryos showed equal *tbx-38* expression in ABal and ABar). (D″) *nos-2* (P3-specific) marks the posterior of the embryo. (E) AB descendants from six replicates of the 16-cell stage embryo. (F) Genome browser tracks of ABxxx transcriptomes, sorted into four groups based on a PCA using only notch target gene expression (shown in Figure S2D). Last exons only are shown. (G) Example of smFISH targeting *hlh-27* (G′, yellow arrows) and *ref-1* (G″, purple arrows) transcripts in intact 15-cell stage embryos (both patterns seen in 100% of embryos, *hlh-27* n=5, *ref-1* n=2). (H) Example of smFISH targeting *tbx-38* (H′, yellow arrows) in intact 15-cell stage embryos (pattern seen in 100% of embryos, n=14). *nos-2* (P_4_-specific) marks the posterior of the embryo.

To filter for informative genes to use in our PCAs, we designed an algorithm to select genes that are reproducibly differentially enriched between cells of the embryo (details in Experimental Procedures). To group replicates of each cell type together, we performed a PCA on all transcriptomes of a given stage using just those filtered genes. We inspected plots of the first and second principle components for distinct groups consisting of one transcriptome from each embryo, which suggest grouping by shared cell-specific features (Figure 2B,E,I,N). We interpreted a group with exactly one cell from each embryo as comprising the replicates of a single (albeit unknown) cell type.

Each PCA tended to isolate only the most dramatically distinct cell types (Figure 2E,I,N). To then identify cell types with more subtle distinguishing features, we removed the transcriptomes that had already clustered out into independent groups, and re-ran the gene selection algorithm and PCA with just the remaining cells. In this way, we continuously enhanced our resolution and split groups of cells off based on increasingly subtle differences (Figure 2F,J,K,O,P; arrows show the cluster of remaining transcriptomes that were put through the next PCA iteration). We chose this iterative PCA approach because it allowed us to take advantage of a unique feature of the *C. elegans* embryo: Each embryo sampled from a given stage generated an identical number of transcriptomes, representing exactly the same set of cell types. Many transcriptome clustering methods define clusters of unspecified size (Yan et al. 2013; Jaitin et al. 2014; Grün et al. 2015; Zeisel et al. 2015), but for this experiment it was most informative to identify groups consisting of exactly one transcriptome from each embryo (see Discussion). The simplest way to achieve this was to inspect the results of a PCA plot for isolated groups of transcriptomes that consisted of one transcriptome from each replicate (Figure 2).

The cells of the 2-cell stage embryo (AB and P_1_) have noticeably different sizes, which allowed us to identify these cells during sample collection. We were able to use this previous knowledge to test the accuracy of our gene selection algorithm and PCA approach. We found that our strategy did in fact allow us to independently and accurately distinguish between these two cell types; all AB cells fell on one side of the first principle component, while all P_1_ cells fell on the other side of the principle component (Figure 2B). The germ cell precursors in subsequent stages (P_2_ at the 4-cell stage and P_3_ at the 8-cell stage) were noticeably smaller than the others and so were also identified upon collection. These cells successfully segregated from the other cells types by our algorithm and PCA (Figure 2E,I). The independent identification of these cells as replicates of each other further validated our algorithm as an effective unsupervised method for selecting informative genes.

To assign identities to groups of cells distinguished by PCA, we examined genes that are known to be expressed in specific cell types. For example, *med-2* is known to be expressed in EMS at the four-cell stage (Maduro et al. 2001). Our transcriptome data shows high *med-2* levels exclusively and robustly in one distinct group of replicates at the four-cell stage (Figure 2G). Based on these observations, we concluded that this cluster consists of the EMS transcriptomes. Similarly, using known markers of cell identity, we verified AB and P_1_ cells at the 2-cell stage, AB daughter cells (ABa and ABp, referred to collectively here as ABx) and P_2_ cells at the 4-cell stage, MS, E, C, P_3_ and AB granddaughter cells (ABxx) at the 8-cell stage, and MS daughters (MSx), E daughters (Ex), C daughters (Cx), D, P_4_, and AB great-granddaughters (ABxxx) at the 16-cell stage (Figure 2, Extended Methods).

### Anterior Cells of the Embryo were Indistinguishable from Each Other by an Unsupervised Multi-Gene Approach, but Show Differential Enrichment of Notch Target Gene mRNAs

For both the 8-cell and 16-cell stages, our PCA approach did not visibly distinguish the descendants of AB from each other (Figure 2K). These results indicate that the transcriptomes of AB descendants at these stages were very similar to each other. This is consistent with the fact that very few genes are known to be differentially expressed between these cells (Priess 2005).

To distinguish between these transcriptomes we examined them for transcripts of a few genes whose proteins are known to be differentially expressed between these cells, namely members of the notch signaling pathway, *hlh-27*, *ref-1* and *tbx-38* (Neves & Priess 2005). We queried all transcriptomes of the AB descendants at the 8‐ or 16-cell stages for transcripts of these three genes, and found that they offered enough information to partition these transcriptomes into four cell types at the 8-cell stage and four pairs at the 16-cell stage (Figure 3B,F, Extended Methods).

To match each hand-sorted group of transcriptomes to a specific cell identity, we performed single molecule FISH (smFISH) on these notch targets in intact 8- and 16-cell embryos. We analyzed micrographs to determine which cell of the embryo expressed each of the distinct combinations of notch targets seen in our data. At the 8-cell stage *hlh-27* transcripts were the most highly enriched in the ABpl and ABpr cells, *ref-1* transcripts were enriched in ABpl cells, and *tbx-38* transcripts were detected at very low levels primarily in ABal (Figure 3C,D). At the 16-cell stage *hlh-27* was enriched in all AB descendants except the ABalx (ABala and ABalp) cells, *ref-1* was detected in ABarx and ABprx cells, and *tbx-38* was detected in ABalx and ABarx cells (Figure 3G,H). This smFISH data in combination with the scRNA-seq data for these notch targets allowed us to sort and identify transcriptomes into the four cell types at the 8-cell stage (ABal, ABar, ABpl and ABpr), and into four pairs of cell types at the 16-cell stage (ABalx, ABarx, ABplx and ABprx) (Extended Methods).

The notch targets mentioned above are critical for cell fate specification of these anterior cells, and are activated via signaling from neighboring cells (Priess 2005). A previous study that sequenced all AB descendants together after allowing them to grow outside of their native embryonic environment showed no *hlh-27* expression in these cells, suggesting that key fate-determining signaling events may have been prevented (Hashimshony et al. 2015). This indicates that processing cells around 10 minutes after dissection, as we did, produces results that more accurately reflect the biology of intact embryos.

Together, our data reveal that the transcriptomes of AB descendants are almost indistinguishable from one another except for transcripts of a few genes, whereas P_1_ descendants show hundreds of differences from one another. Some pairs of cells were ultimately indistinguishable from each other by our method, but because we sequenced each cell independently, this decreased resolution is a reflection of the biology of the cells.

### The Transcriptional Lineage Expands Upon Known Gene Expression Patterns During Development and Increases Their Resolution

Having assigned cell identities to each transcriptome in our dataset, we first confirmed that the data and our identity assignments reflected certain known expression patterns. We queried our dataset for expression patterns of *sdz-38* (which encodes a putative zinc finger protein that is expressed in the MS cell; Robertson et al. 2004), *tbx-37* (a T-box transcription factor found in ABa descendants; Neves & Priess 2005), *ceh-51* (a homeodomain protein expressed in the MS lineage; Broitman-Maduro et al. 2009), *elt-7* (a GATA-type transcription factor that induces gut specification in the E descendants; Sommermann et al. 2010), *cwn-1* (a wnt ligand that is expressed in the C and D cells; Gleason et al. 2006), and *cey-2* (a putative RNA binding factor restricted to the germ line; Seydoux & Fire 1994). None of these genes were used to previously identify each cell type (Figures 2 and 3), allowing us to use their expression patterns to independently test the validity of our data and cell assignments. Our scRNA-seq data reflect the expected patterns for all six of these genes (Figure 4A, key in Figure 1F), and in addition quantified their expression in each cell, along with 8,749 additional detected genes (Figure 4B).

**Figure 4:**
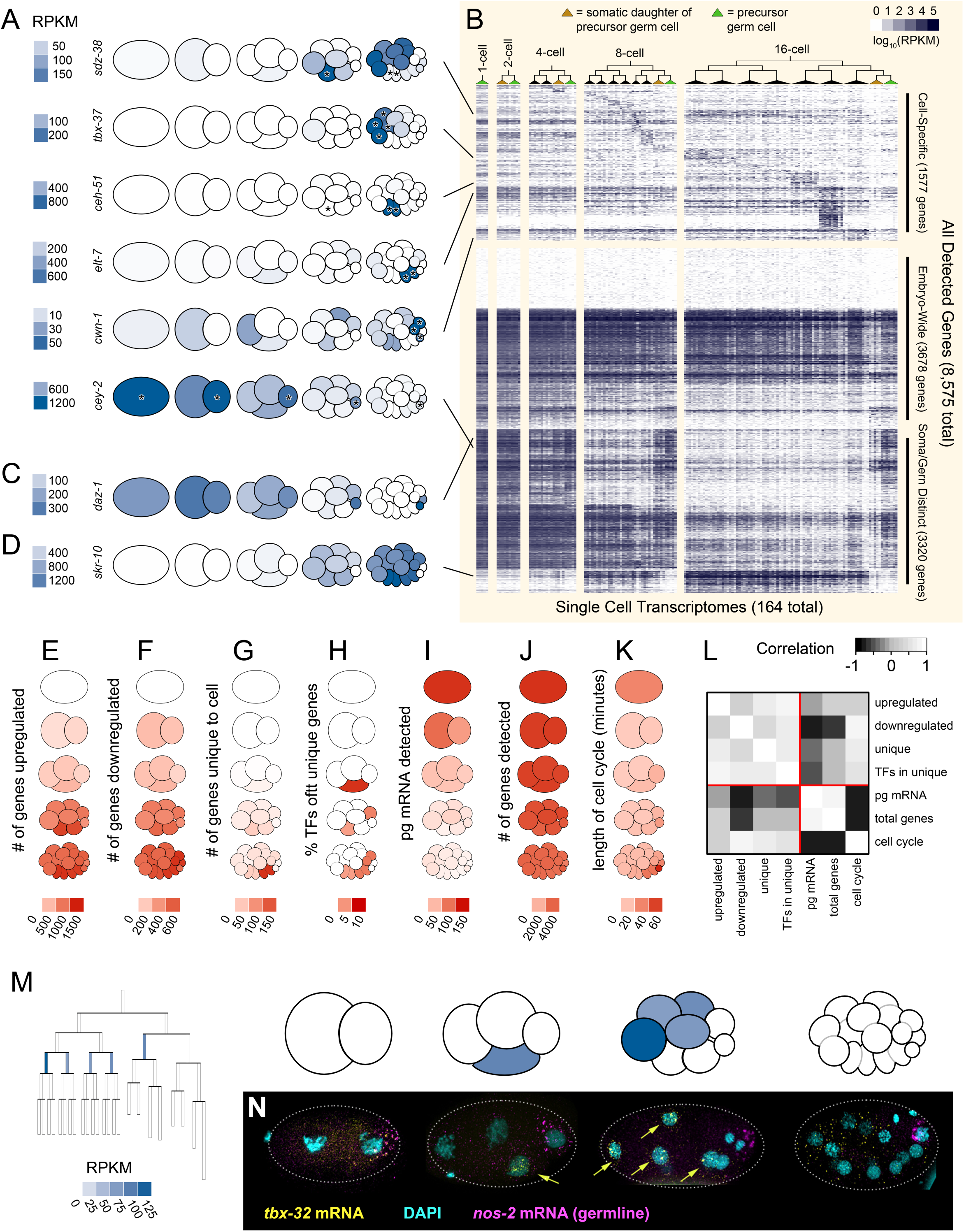
Differential activation of the zygotic genome in each cell lineage. (A) Transcript abundances of six genes with previously known expression patterns, heat-mapped on to pictograms of the embryo (key in Figure 1F). Asterisks indicate the cells in which we expected expression, based on the literature; *sdz-38* expected in E, Ex (Ea and Ep); *tbx-37* expected in ABalx (ABala and ABalp), ABarx (ABara and ABarp); *ceh-51* expected in MS, MSx (MSa and MSp); *elt-7* expected in Ex (Ea and Ep); *cwn-1* expected in Cx (Ca and Cp), D; *cey-2* expected in P_0_, P_1_, P_2_, P_3_, P_4_ (references in Main Text). (B) Heatmap of transcript abundances of all 8,575 present genes (y-axis) in each cell throughout time and space (x-axis). Only transcriptomes that passed quality filtration were plotted (164 out of 219). The y-axis along the top third of the heatmap is scaled twice as large as the bottom two thirds, to show detail. (C) Transcript abundance data for *daz-1* (a maternally inherited gene required for meiosis; Karashima et al. 2000), an example of a transcript we detected in only the germ cells and their sister cells. (D) Transcript abundance data for *skr-10* (a member of the ubiquitin ligase complex; Yamanaka et al. 2002), an example of a transcript we detected in only somatic cells. (E) The number of upregulated genes for each cell type. Genes were scored as upregulated in a cell if their transcripts were at least twice as abundant as in any ancestors of that cell‥ (F) The number of downregulated genes for each cell type. Genes were scores as downregulated in a cell if their transcript abundance was half or less that of an ancestor. (G) The number of cell-specific, or unique, genes. Genes were scored as unique to a cell type if their transcript abundance was at least 10 times higher than in any other cell type in the dataset. (H) Percentage of each cell type’s unique genes (G) that are transcription factors. (I) Mass of mRNA per cell as calculated using concentrations of control mRNA spike-ins. (J) Number of genes detected above 25 RPKM in each cell. (K) Length of cell cycle for each cell. (L) Pearson correlation of E-K across all cell types (excluding germ cell precursors, which are known to be transcriptionally distinct; Schaner & Kelly 2006). (M) A cell lineage map and a pictogram of the 1-through 16-cell stages. Color corresponds to transcript abundance data for *tbx-32* in each sample. (N) smFISH of *tbx-32*. 100% of 2-cell stage embryos (n=2), 83% of 4-cell stage embryos (n=6, one embryo showed ubiquitous staining), 100% of 8-cell stage embryos (n=6), and 100% of 16-cell stage embryos (n=3) showed this pattern.

Low-input transcriptomes for some of these cells, including AB and P_1_, have previously been generated (Hashimshony et al. 2012; Osborne Nishimura et al. 2015; Hashimshony et al. 2015), as have whole-embryo microarray timecourses of *C. elegans* development (Baugh et al. 2003; Baugh et al. 2005). We compared our scRNA-seq data to data from two previous studies (Hashimshony et al. 2015, Osborne Nishimura et al. 2015) that each used different methods to sequence mRNA from AB and P_1_ cells (2-cell stage). We calculated enrichment index values for each gene (a product of the gene’s AB/P_1_ fold change and the gene’s average expression, Experimental Procedures). To measure the agreement between each study, we compared the enrichment index values calculated from each study’s data. All studies were positively correlated with one another to similar extents, and the correlation increased when only significantly differentially enriched genes were compared (Figure S3A).

We analyzed data from the 2005 whole-embryo microarray timecourse (Baugh et al. 2005) to test if our low-input transcriptomes reflect the patterns identified by a higher-input but lower-sensitivity experiment. We searched for genes whose transcript levels either increased or decreased by twofold over time in the microarray data and identified 1,935 and 2,164 genes respectively. Transcripts whose levels increased or decreased in our dataset included 91% and 97% of those detected in the earlier dataset. In addition, we identified 7,763 other transcripts that increased or decreased over time, many of which had very low expression levels, presumably undetectable by microarray, and 1,053 for which there was no microarray probe in the previous experiment (Figure S3B). This result suggests that even though the transcriptomes we present here were generated from just picograms of mRNA, they capture the patterns described by a higher-input method, but with much greater sensitivity and resolution.

### Transcriptional Dichotomy Between Germ Cells and the Soma

Visualization of transcript levels for all 8,575 genes detected across all cell types revealed three distinct trends of gene expression (Fig. 4B): First were transcripts absent in the zygote and only detected in subsets of cell types, suggesting cell-specific transcription (Fig 4B top). Second were transcripts present in the zygote that then became detected at lower levels over time in an embryo-wide fashion, suggesting global mRNA degradation (Fig. 4B center). Third were transcripts that were differentially abundant between somatic cells and germ cell precursors (Fig 4B bottom). Within this third group, in some cases transcripts became undetectable over time in the somatic cells but remained detectable in the germ cell and the immediate sister of the germ cell (as in *daz-1*, a gene required for oogenesis; Karashima et al. 2000, Figure 4C). In other cases, genes became detectable over time in the somatic cells, while remaining undetectable in germ cells and their sisters (as in *skr-10*, a core component of the ubiquitin-ligase complex; Yamanaka et al. 2002; Figure 4D). Transcriptional quiescence in the germline is a feature that many organisms share (Deshpande et al. 2004; Cheung & Rando 2013). Our dataset quantifies this phenomenon and identifies the genes affected by it.

### Differential Activation of the Zygotic Genome Among Cell Lineages

Fundamental events of embryonic development start earlier in the *C. elegans* embryo than in many other model organisms. Cell-fate determining steps begin as early as the 2-cell stage, and gastrulation begins at the 26-cell stage. Within the embryo, certain cells engage in these events earlier than others. For example, gastrulation begins earliest in the E descendants and follows later in other cells (Nance et al. 2005). By further example, at the 16-cell stage the P_1_ descendants (which we will refer to as posterior cells) include 4 cells that are already restricted to a single fate, while none of the AB descendants (which we will refer to as anterior cells) are as fully fate-restricted (Figure 1A). Based on this, we hypothesized that transcriptomes change more dramatically in the more fate-restricted posterior cells than the anterior cells. To quantify the extent to which transcriptomes of each lineage change over time, we asked how many genes were detected as having increased or decreased transcript levels in each cell when compared to the cell’s parent. We found a higher number of both increasing and decreasing transcript levels in the non-germ descendants of the P_1_ cell (Figure 4E,F) than in AB descendants, supporting our hypothesis that there is more dynamic gene regulation in these cells than in the AB descendants. We wondered whether this apparent increased dynamism of gene regulation (number of transcripts increasing or decreasing in abundance) in the non-germ posterior cells could be related to other feature of these cells, such as greater mass of mRNA (Figure 4I), greater transcriptome complexity (Figure 4J), or longer cell cycle (Figure 4K; Wormbase 2007). By the 16-cell stage, the total mass of mRNA and the number of detected transcripts in each cell negatively correlated with the dynamism of gene regulation in the posterior cells (average R = −0.52 and −0.19) while the length of the cell cycle positively correlated with the dynamism of gene regulation (average R = 0.51, Figure 4L). This suggests to us that there are cellular features broadly associated with a cell lineage’s progression through the maternal to zygotic transition, including fewer total transcripts and a longer cell cycle.

To quantify the extent to which each cell’s transcriptome is unique, we evaluated the number of genes with transcripts detected exclusively in that cell and no others. Again we saw higher numbers of unique transcripts in the non-germ descendants of the P_1_ cell. The cell type with the highest number of uniquely expressed genes (176) was the Ex cells (Ea and Ep; Figure 4G). These cells have already established an endoderm-specific transcription program (Maduro 2010), and are minutes away from initiating gastrulation by moving from the outside of the embryo to the inside (Nance et al. 2005). These are also the first cells that have a gap phase in their cell cycles, taking 40 minutes to divide compared to ~20 minutes in the other cells of this stage (Edgar & McGhee 1988). Because many of the posterior cells become restricted to a single fate before the anterior cells do, we hypothesized that the posterior cells might express a greater number of cell-specific transcription factors. For each cell type, we calculated the percentage of that cell’s unique genes that were transcription factors. We found a larger proportion of mRNAs encoding transcription factors uniquely in the posterior cells (Figure 4H), suggesting that these cells are initiating lineage-specific transcriptional programs.

### Genes with Spatially Dynamic Expression

When a given transcript is detected across multiple temporal stages in an embryo, the most parsimonious explanation is that the transcript is inherited from parent cells to daughter cells lineally. While we expect some genes to contradict this assumption and be uniquely expressed in cells that are not related by lineage, such a scenario cannot be detected with a whole-embryo timecourse. The present dataset has a high enough resolution both temporally and spatially that we were able to identify transcripts whose overall expression is continuous throughout consecutive stages, but that are detected in different cell lineages throughout those stages. One such example is *tbx-32* (Figure 4M), which was robustly detected in EMS at the 4cell stage but absent in the daughters of EMS (E and MS) at the following stage. Instead *tbx-32* transcripts appeared in anterior cells ABal, ABar, ABpl and ABpr (also referred to here as ABxx), that are not directly related to EMS by lineage. We found five more genes (*tbx-31*, *tbx-40*, Y43D4A.6, Y116A8C.20, ZK666.1; Figure S5A) that have patterns similar to *tbx-32*, suggesting that a common mechanism may be regulating all of these genes.

To test the validity of this cross-lineage expression pattern, we performed smFISH on intact embryos. We detected *tbx-32* transcripts in EMS at the 4-cell stage and in AB descendants at the 8-cell stage, as our RNA-seq data predicted (Figure 4N). *tbx-32* transcripts were more abundant in the 16-cell stage by smFISH than we anticipated from our RNA-seq dataset, but partially degraded transcripts may be more detectable by smFISH (which recognizes many sequences in the transcript) than by the RNA-seq method we used (which requires the presence of a polyadenylated tail for detection). This smFISH data allowed us to describe the *tbx-32* expression pattern with an even higher temporal resolution than in the transcriptional lineage (Figure S5B). The smFISH data revealed nuclear localization of *tbx-32* transcripts early in the EMS and ABxx cell cycles, and cytoplasmic localization later in these cell cycles. This sequence of localizations suggests that the dynamic pattern of *tbx-32* expression is due to zygotic transcription in these cells.

### scRNA-seq Reveals Genes that are Required in Combination for Embryonic Development

Most of our understanding of gene functions in the early *C. elegans* embryo has been gleaned from genetic screens that investigate phenotypes of single-gene disruptions. A blind spot of genetic screens, though, is genes with partially redundant functions. These genes yield modest or no detectable phenotype when disrupted because of other genes that are capable of fulfilling at least part of their function. An estimated 32% of *C. elegans* genes have one or more paralog (Woollard 2005), a common feature of partially redundant genes. We were curious to see whether our dataset could provide insight into genes that are necessary for development, but whose importance has eluded discovery because of their overlapping function with other genes. We searched our data for groups of genes that meet two criteria; the genes in the group must be synexpressed (having transcripts that are similarly enriched in the same cell type; Niehrs & Pollet 1999) and must be similar to one another in sequence.

We found 99 sets of genes that were synexpressed and paralogous (Experimental Procedures), making them candidates for sets of partially redundant genes (Figure 5A, Figure S3). As a control, we scrambled the gene names in our dataset 100 times and found an average of 14 synexpressed paralogous gene sets in these permutations (Figure 5B). The 99 true sets consisted of 2-7 genes each, over half of which have no known or predicted function (WormMine 2016).

**Figure 5:**
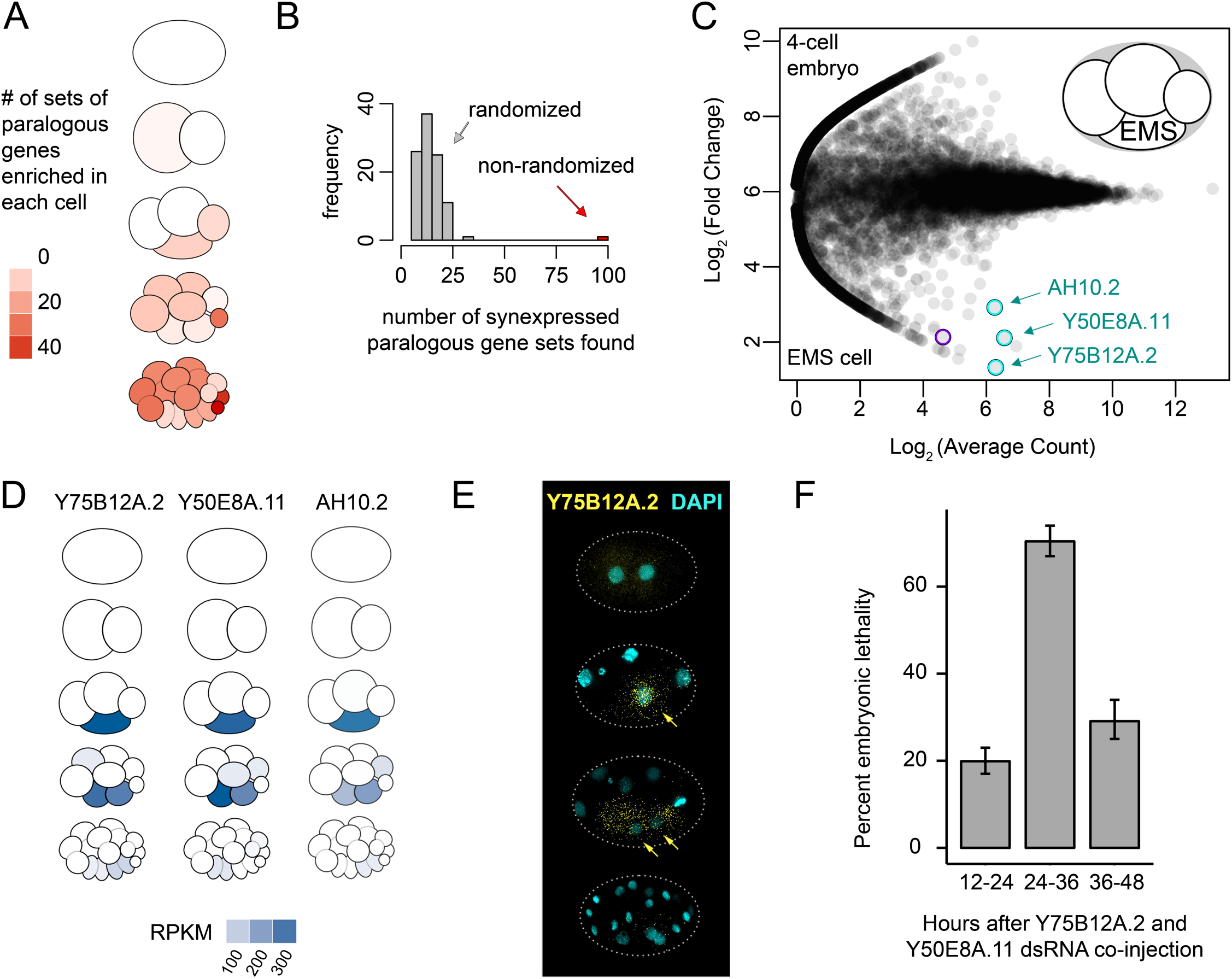
Paralogous, synexpressed genes are critical for development in a partially redundant manner. (A) Heatmap of the number of synexpressed paralogous gene sets identified in each cell type. (B) Histogram of the number of synexpressed paralogous gene sets detected in our dataset (red bar) or in 100 datasets randomized by scrambling gene names without replacement (gray bars). (C) MA plot comparing gene expression in the EMS cell to the other cells of the 4-cell stage. Purple circle highlights *med-2*, an example of a known gene with EMS-enriched transcripts. Cyan circles highlight three paralogous genes that have no phenotype when knocked down individually. (D) Pictograms showing quantitative transcript abundance data for the three genes identified in (C). (E) smFISH of Y75B12A.2 (pattern seen in 100% of embryos, n=4 for 2-cell stage, n=6 for 4-cell stage, n=4 for 8-cell stage, n=3 for 16-cell stage). (F) Lethality phenotype observed in embryos in which Y75B12A.2 and Y50E8A.11 were targeted by co-injection of dsRNA.

Of the 99 sets we identified, one set included three genes (Y75B12A.2, Y50E8A.11 and AH10.2) that were specifically enriched in the endomesoderm precursor, EMS, at the 4-cell stage (Figure 5C). These three transcripts were not detected before the 4-cell stage, but once they appeared, their transcript levels were high and their expression patterns were transient and cell-specific (Figure 5D). We validated this pattern for one gene of this set, Y75B12A.2, via smFISH (Figure 5E).

Knocking down each of these three genes independently by RNAi in previous studies resulted in no detectable phenotype (Kamath et al. 2003; Sönnichsen et al. 2005). Acting on our hypothesis that these genes have partially overlapping functions, we knocked down two of them (Y75B12A.2 and Y50E8A.11) by co-injection of double stranded RNAs. We found that mothers in which both Y75B12A.2 and Y50E8A.11 were targeted laid embryos with a 70.1% rate of embryonic lethality (Figure 5F), compared with no phenotype (<10% embryonic lethality) when knocked down independently, as reported in the literature (Kamath et al. 2003; Sönnichsen et al. 2005). This supports our hypothesis that the transcriptional lineage can help identify partially redundant genes that yield phenotypes only when targeted in concert.

### An Interactive Data Visualization Tool to Explore Our Gene Expression Data

To maximize the accessibility of our data, we developed an interactive data visualization tool (available in Chrome and Firefox browsers at http://tintori.bio.unc.edu). With this tool, the user can select which two cells or embryos they wish to compare, and generate a differential gene expression plot that highlights all of the transcripts enriched specifically in either sample (Figure 6). The tool allows hypothesis-driven analyses, in which the user can query known genes of interest, as well as exploratory analyses, in which the user can discover new genes of interest.

**Figure 6:**
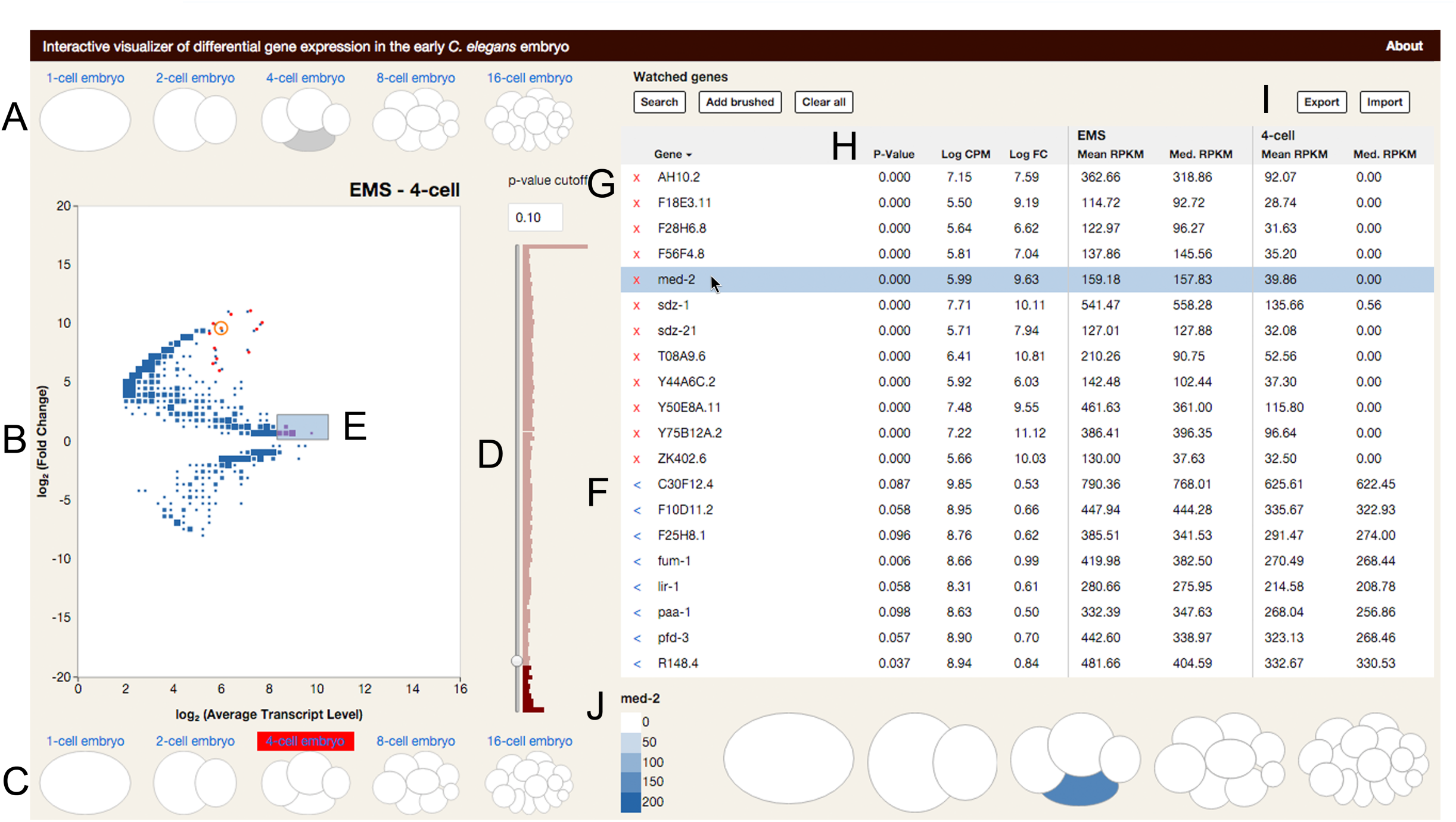
An interactive data visualization tool for querying the transcriptional lineage. Still image of data visualization tool. Full version available in Chrome and Firefox broswers at http://tintori.bio.unc.edu. (A-C) Sample selection. The user clicks on the cells or whole embryos they wish to compare on the top (A) and bottom (C) of the plot. When a new sample is selected, the plot (B) is drawn to reflect the selected comparison. Size of points in B scales to the number of genes represented by each dot. (D-H) Gene selection. (D) The user can filter genes by P-value of differential enrichment between samples. (E) Clicking on a point or selecting a swath of points on the plot adds genes and their data to the Selected Genes table (F). Known genes can be added directly, by typing their names into the search bar. (G) The Watched Genes table is curated by adding Selected Genes individually or in bulk. (H) The Watched Genes table can be exported, and lists of genes can be imported to the Watched Genes table in bulk. (I-J) Gene expression metrics. (I) The gene tables are sortable by name, average expression level, fold change, significance of differential enrichment, and expression levels in either sample being compared. (J) Clicking on a gene in the table reveals a cartoon of the embryo over all five stages. Each cell is colored corresponding to the transcript level of the highlighted gene.

Our scRNA-seq data may be used to explore many fundamental aspects of development, such as specification of distinct cell types such as muscle or intestine, and cell behaviors such as cell cycle control or morphogenesis. We hope our visualization tool will invite researchers working on these topics to explore our dataset.

## DISCUSSION

### A transcriptional lineage to complement the completely defined cell lineage of the *C. elegans* embryo

For decades the *C. elegans* embryo has been a powerful tool for studying cell biology and development, largely because of its invariant cell lineage (Sulston et al. 1983). Here we present a transcriptional lineage that, when paired with the cell lineage, describes the suite of transcripts present in early embryonic cells as they diverge in fate, morphology, and behavior. As technology improves, scRNA-seq of cells beyond the 16-cell stage will become possible, ideally allowing the possibility of a transcriptional map for every cell at every stage of development. The challenge of *post hoc* cell identification, explored in a previous study (Hashimshony et al. 2012) and in this manuscript, will continue to be relevant at these later stages of development.

Several research groups have previously performed scRNA-seq on human and mouse cells, and identified their cell types *post hoc* by the transcriptomes (Grün et al. 2015; Yan et al. 2013; Biase et al. 2014; Xue et al. 2013; Zeisel et al. 2015; Jaitin et al. 2014; Trapnell et al. 2014; Satija et al. 2015; Achim et al. 2015; Pollen et al. 2014). The invariant development of *C. elegans* provides a constraint on the possible identities of each transcriptome. This advantage can help guide cell-type identification and is not present in other systems. For example, because each 4-cell embryo yields exactly 1 transcriptome each of exactly 4 cell types, we know that out of our total of 20 unidentified transcriptomes from this stage (4 cells x 5 replicates), exactly 5 of them are from P_2_ cells, 5 are from EMS cells, 5 are from ABa cells, and 5 are from ABp cells. Because we know that a cell from one embryo will have exactly one counterpart in each other embryo, we were able to plot Principle Component Analyses of all transcriptomes, as in Figure 2E,F, and look for clusters containing one cell from each embryo. Given the unique constraints of this system, such clustering suggests that all replicates of a single cell-type are grouping together. The fact that known cell-type markers show transcript enrichment patterns that are consistent with our replicate grouping indicates the accuracy of our gene filtration and iterative PCA approach (Figure 2C,G,LQ).

Another difference between our study and previous scRNA-seq studies that identified cell types *post hoc* is that the *C. elegans* cells used in the present study divide about every 20 minutes, whereas the human and mouse cells of previous studies divide approximately every 24 hours. Given this comparatively short cell cycle, it is remarkable that the posterior *C. elegans* cells have such distinct transcript signatures, and by the same token perhaps not surprising that the anterior cells are difficult to distinguish.

### Cells of the Early Embryo Can Be Identified by Their Transcriptomes Alone

We have assigned a cell identity to each transcriptome based on its transcript abundance data, cross-referenced to known expression patterns and *in situ* RNA hybridization. The transcriptomes of some cell types (particularly the P_1_ descendants) grouped together tightly, were clearly distinct from other cell types, and had identities confirmed by well-studied genes, making us confident in our assessments. For the anterior cell types whose transcriptomes were less distinct from one another, we have a lower confidence in our assignments (as in Figure 3D′). We consider this paucity of distinguishing features to be an interesting biological result, suggesting that it would make little difference if transcriptomes were mis-assigned between these cell-types. The current understanding of these cells’ developmental potential supports the notion that they should be difficult to distinguish from each other. For example, the sister cells ABa and ABp of the 4-cell stage are initially developmentally equivalent, and the differences between them are not established until after cytokinesis separates them (Priess & Thomson 1987). In the future, if features are identified that more clearly differentiate these cell types, our existing single-cell transcriptomes can be revisited with those features in mind.

Previous studies that have measured transcript abundance in cells of the early embryo have either measured whole-embryo transcript levels (Baugh et al. 2003; Baugh et al. 2005; Levin et al. 2012), or measured only parts of the embryo at a single-cell resolution and the rest of the embryo in clusters of related cells (Hashimshony et al. 2015). These clusters of cells were sampled by dissecting embryos starting at the 2-cell stage and allowing the isolated cells to divide in culture, then sequencing the group of descendants. This allowed descendants of founder cells to be harvested at later time points than in our study, but kept the cells naïve to critical signaling events that take place in intact embryos. With our dataset, by leaving all cells intact in the embryo until minutes before sampling, we captured single-cell transcriptomes while allowing the cell-cell signaling necessary for proper development to occur, and we detected the transcriptional results of this signaling (Figure 3B,F).

### A Stark Contrast in mRNA Composition Between Germ Cell Precursors and Somatic Cells

One pattern that is apparent when comparing gene expression across all cell types (Figure 4B) is that there is a prevalent distinction between the mRNA composition of the somatic cells and the germ cells (including the somatic sister of each germ cell precursor). Previous studies, such as Seydoux & Fire 1994, have observed this contrast in transcript composition between the germ and soma. Their reliance on *in situ* hybridization necessarily restricted the number of such genes they were able to study (10 genes), whereas the present genome-wide study expands their findings to thousands of genes. Differences between “immortal” germ cells and “mortal” somatic cells have fascinated researchers for over a century (Weismann 1893; Boveri 1910; Schierenberg & Strome 1992; Lai & King 2013; Lehmann & Ephrussi 2007; Yamanaka 2007). The present dataset quantifies this dichotomy and the transition from one state to the other over time. This dataset includes before, during, and after snapshots of somatic descendants of germ cell precursors, in their transition from the germ-like profiles of their parent cell to the somatic profiles of their descendants.

### Cross-lineage Expression Patterns Highlight Genes that May Share Mechanisms of Gene Regulation

*tbx-32* and the five other genes with similar expression patterns are examples of genes whose expression is not continuous from parent to daughter cell, but rather appears in one cell type (EMS) at one stage, then in a different lineage of cells (ABxx) at the next stage. The EMS cell at the 4-cell stage and one of these ABxx cells (ABar) at the 8-cell stage have another feature in common, which is that both orient their mitotic spindles in response to Wnt signaling (Walston et al. 2004).The fact that this specific expression pattern is shared by several genes suggests that a common mechanism may be regulating all of these genes, possibly the previously characterized Wnt signaling. Alternatively, these six genes may play a role in establishing which cells are capable of responding to Wnt signaling.

### Identifying Partially Redundant Critical Regulators of Development

One of our initial hypotheses was that many genes regulating development are partially redundant with other genes, making them elusive to genetic screens but detectable by RNA-seq. We identified 99 sets of gene paralogs whose similar expression patterns suggest that they may have partially redundant functions. We investigated one such set of genes and found a high level of embryonic lethality when multiple genes were targeted together, even though each gene has been reported to have no phenotype when knocked down individually. Thus, our dataset may be well-suited to identify additional regulators of development that have not previously been detected.

## EXPERIMENTAL PROCEDURES

### Worm husbandry and embryo dissections

All worms were grown at 20°C and dissected at room temperature (21-24°C). Single embryos were selected at 10-20 minutes before the desired stage, eggshells were weakened with 10% bleach in Egg Buffer for 3 minutes followed by a 3-5 minute incubation in 20 units/mL chitinase and 10 mg/mL chymotrypsin in Egg Buffer (modified from Edgar & Goldstein 2012). Cells were manually teased apart by aspiration in Shelton’s media. Cells used for 16-cell samples were separated from each other at the 12 cell stage, so that final cell divisions could be observed immediately before embryos were collected into 16 tubes at the 24-cell stage (all 16 AB cells were collected as 8 pairs of sister cells). All cells from an embryo were simultaneously flash frozen in liquid nitrogen and stored at −80°. Only complete sets of cells were collected for 1-cell through 8-cell samples. For 16-cell samples, sets of cells were collected if at least 14 cells remained intact. Of the 16-cell replicates that passed quality filtration, two replicates had all 16 samples, three replicates were missing one descendant of P_1_, and one replicate was missing 2 descendants of AB. All replicates of any given stage were collected on independent days, and libraries were prepared on independent days, with the exception of the first and second 1-cell stage replicates.

### RNA preparation, sequencing, and RPKM generation

cDNA was generated using the SMARTer Ultra Low RNA Input for Illumina Sequencing Kit, and sequencing libraries were prepared using the Nextera XT kit, both according to manufacturers instructions. ERCC RNA spike-in controls were added at a 1:2,000,000 dilution, with the exception of the first replicate of the 4-cell stage, which was diluted 1:500,000. All libraries were sequenced on a Hi-Seq 2500 Illumina machine, for 50 cycles from a single end. Sequences from this study are available at NCBI GEO GSE77944 (reviewer access: http://www.ncbi.nlm.nih.gov/geo/query/acc.cgi?token=uhqtwmumzvshroi©acc=GSE77944). Identical reads were collapsed using fastx_collapser (FASTX Toolkit 0.0.13.2; Gordon & Hannon 2010) and aligned to the ce10 genome using Tophat2 (2.0.11; Kim et al. 2013). Count files were generated using HTseq-count (HTSeq 0.5.3p3; Anders et al. 2014) and a WBcel235.78.gtf reference file (ftp.ensemble.org).

### Transcriptome analyses

Analyses were performed in R. All scripts and functions are available at https://github.com/tintori/transcriptional-lineage. In all analyses, genes were only considered expressed if their transcripts were detected above 25 RPKM.

### RNAi

Double stranded RNAs for Y75B12A.2, Y50E8A.11 and *par-6* (negative control) were made using primers with the following sequences: ATGTGCTCTTATTCTCTTGCTCCGGC, TCAAAAAAGGCTATGTGGCTCTGCGG, ATGGTTTCTGCCACTGGACATCTTCG, AATTTATTCATTTAGTTATATTTTTT, ATGTCCTACAACGGCTCC and TCAGTCCTCTCCACTGTCC. dsRNAs were generated with Promega’s T7 Ribomax Express RNAi kit, according to manufacturers instructions. RNAs were combined and diluted to a total of 1 ug/uL for each condition. 20-30 young adult worms were injected for each condition. After injection, worms were moved to fresh plates every 12 hours, and plates were assayed 24 hours later for number of larval offspring and unhatched eggs. Embryonic lethality was calculated as the percent of unhatched embryos remaining 24 hours after worms were removed from the plate out of the initial number of unhatched embryos upon removing mothers.

### Single molecule fluorescent in situ hybridization

Stellaris smFISH fluorescent probe sets (Biosearch) were generated to target transcripts. N2 worms were grown at 20°C to gravidity, bleached for embryos, resuspended in −20°C methanol, freeze cracked in liquid nitrogen, and fixed at −20°C for 24–48 hours ( as in Shaffer et al. 2013). Embryos were equilibrated in WB1 (100 mg/ml dextran sulfate, 10% formamide, 2 x SSC), hybridized in hybridization buffer (100 mg/ml dextran sulfate, 1 mg/ml *E. coli* tRNA, 2 mM vanadyl ribonucleoside complex, 0.2 mg/ml BSA, 10% formamide) containing 50 picomoles of each primer set (Ji & van Oudenaarden 2012). Hybridization at 30°C overnight was followed by two 10 minute WB1 washes and one 10 minutes WB1/DAPI staining at 30°C. This was followed by three 2 x SSC washes at room temperature. Embryos were mounted as described by Ji & van Oudenaarden 2012, using SlowFade (Life Technologies) to prevent photobleaching. smFISH images were acquired using a Photometrics Cool Snap HQ2 camera on a DeltaVision-modified inverted microscope (IX71; Olympus), with a UPlanSApo 100 x (1.40 NA) objective and SoftWorx software (Applied Precision) using fixed exposure and acquisition conditions (0.3 μm z-stacks, up to three wavelengths using AlexaFluor emission filters). Images were deconvolved using DeltaVision deconvolution.

### Defining paralogous sets of genes

Genes were BLASTed against the *C. elegans* EST collection with a BLAST e-value threshold of less than 10^-15^. The cut-off of e=10^-15^ was chosen based on *end-1* and *end-3*, a known example of paralogous genes that overlap in function. e=10^-15^ was the most conservative cut off that resulted in *end-1* and *end-3* appearing in each others’ list of BLAST hits.

## AUTHOR CONTRIBUTIONS

Experiments were conceived of by SCT, JDL and BG.

smFISH was performed by EON, all other experiments and analyses were performed by SCT. Data visualization tool conceived of by SCT and PG, and coded by PG.

Manuscript written by SCT.

## ACKNOWLEDGEMENTS

The authors would like to thank Corbin Jones and the UNC High Throughput Sequencing Facility for their generosity during scRNA-seq protocol optimization. The authors declare no conflicts of interest. BG and SCT were supported by NIH grant R01 GM083071. JDL, EON and SCT were supported by NIH grant R01 GM104050. SCT was supported by NIH grant F31 HD088128 and by a NSF GRFP award.

